# Identifying Genes with Location Dependent Noise Variance in Spatial Transcriptomics Data

**DOI:** 10.1101/2022.09.25.509381

**Authors:** Mohammed Abid Abrar, M. Kaykobad, M. Saifur Rahman, Md. Abul Hassan Samee

## Abstract

Spatial transcriptomics (ST) holds the promise to identify the existence and extent of spatial variation of gene expression in complex tissues. Such analyses could help identify gene expression signatures that distinguish between healthy and disease samples. Existing tools to detect spatially variable genes assume a constant noise variance across location. This assumption might miss important biological signals when the variance could change across spatial locations, e.g., in the tumor microenvironment. In this paper, we propose *NoVaTeST*, a framework to identify genes with location-dependent noise variance in ST data. NoVaTeST can model gene expression as a function of spatial location with a spatially variable noise. We then compare the model to one with constant noise to detect genes that show significant spatial variation in noise. Our results show genes detected by NoVaTeST provide complimentary information to existing tools while providing important biological insights.

## Introduction

Spatial variation in gene expression is strongly linked to tissue function and physiology^1–4^. Although the first demonstrations of this phenomenon came from low-throughput techniques, spatial transcriptomics (ST) now allows transcriptome-scale identification of genes that have location-dependent expression. The commercially available Visium ST^5^, for example, captures the transcriptome of a tissue slice in a spatial grid of spots, each spot containing 5–15 cells on average^6^ (Figure 1(A)).

**Figure 1.**
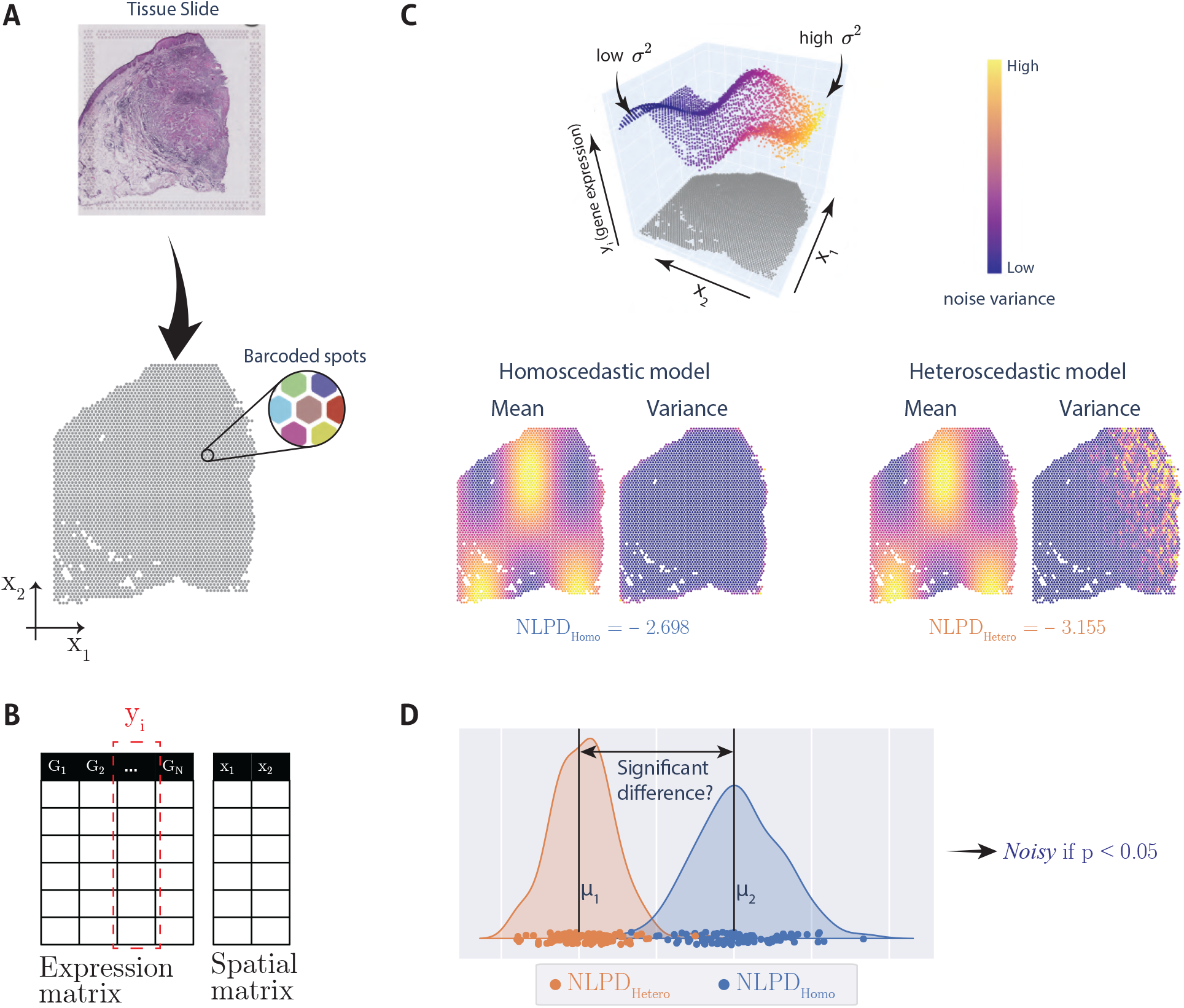
Graphical overview of the NoVaTeST pipeline. (**A**) Visium technology to acquire ST data at pre-designated spatially barcoded spots from a tissue slice placed on top of a slide. (**B**) Representaiton of ST count data. Each column in the Expression matrix represents the expression of a particular gene with spot locations given by the Spatial matrix. (**C**) Simulated gene expression with heteroscedastic noise, along with the two spatial models – homoscedastic and heteroscedastic. The lower NLPD for the heteroscedastic model indicates a better model fitting. (**D**) Comparing the NLPD values of the two models generated from multiple trials to see if there is a statistically significant difference between the two models. A *p*-value (obtained using Wilcoxon signed rank test) less than 0.05 indicates that the heteroscedastic model provides better fit than the homoscedastic model, thus the expression is *noisy*.

Modeling the gene expression as a function of location is the first step toward the spatial variation analysis of ST data. Existing models of these data assume that gene expression noise has a constant variance across locations. Thus, a gene’s expression *y_i_* at location *x_i_* is modeled using a function *f*(*x_i_*) and a Gaussian noise *ε* with mean zero and variance *σ*^2^:

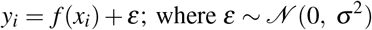

These models have provided valuable biological insights by identifying genes that show significant location-dependent changes in expression^7, 8^. However, we still lack models that assess if the noise variance of a gene’s expression is location-dependent, *i.e*., heteroscedastic^9^. In this case, the variance of *ε* is not an unknown constant *σ*^2^ but a variable 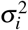 that depends on the location *x_i_*. The premise is set by prior analyses of spatial data in biology^10–12^, economics^13–15^, and robotics^16–18^. For example, the magnitude of imaging noise for apparent diffusion coefficient during whole-body diffusion-weighted MRI is heteroscedastic^10^. Park *et. al*.^12^ used heteroscedastic noise variance to encode confidence levels in predicting the tumor mutation burden from whole slide images. In economics, heteroscedastic regression is used for modeling time-varying volatility and stochastic volatility models in time series data^13–15^. Heteroscedastic regression is also used in robotics^16, 17^, including where multi-modal sensor data are combined for terrain modeling^18^. Other fields where heteroscedastic noise variance is used include biophysical variable estimation^19^, vehicle control^20^, cosmological redshift estimation^21^, etc.

These studies have shown that spatial signals do not always indicate constant variance, and thus suggest modeling *ε* of gene expression with a location-dependent variance 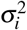. Motivated by this, we propose *<u>no</u>ise <u>va</u>riation <u>te</u>sting in <u>ST</u> data (NoVaTeST)*, a method to identify genes with statistically significant heteroscedasticity. Our method is based on the idea that for genes with location-dependent noise, a heteroscedastic model is expected to yield a better fit compared to a homoscedastic (constant noise) model. One way to measure this “goodness-of-fit” is to use negative log predictive density (NLPD) on a test dataset, “which penalizes over-confident predictions as well as under-confident ones”^22^. As an example, the NLPDs of a homoscedastic model and a heteroscedastic model for a simulated expression are shown in Figure 1(C). Our method compares the NLPDs using rigorous statistical tests to select the best model (homoscedastic vs heteroscedastic) for each gene to find the ones with location-dependent noise. The method also clusters the detected genes with similar noise variance patterns. These gene sets can be used to infer underlying biological signals such as pathway enrichment.

In the context of ST data, spatial variation of noise variance could indicate gene expression variation due to sequencing technology, as well as variation due to biology. For example, a common source of technical noise is the mean-variance relation^23–25^, where the noise variance typically increases with the mean expression of a gene. Several variance-stabilizing transformations are available to remove this type of technical noise^26, 27^. On the other hand, the variation in noise could also be due to underlying biologies, such as cell-type heterogeneity^28^ and phenotypical variation across spatial coordinates. Moreover, if the sample being analyzed is partially affected by some condition, the noise variance for some genes is likely to be different in the affected region compared to the non-affected region^29^.

## Results

### NoVaTeST: A pipeline for detecting genes with spatially variable noise

Briefly, after an initialquality check (QC) filtering, the method first transforms the gene expression count data using the Anscombe technique^26^ as a variance-stabilizing transformation to reduce the effect of the mean-variance relation, followed by regressing out library size effect. Next, for each gene, a regular Gaussian process^30^ (GP) and a heteroscedastic version of the GP^31^ is used to get two models, one homoscedastic and one heteroscedastic, respectively. Finally, statistical model selection technique is used to identify the better fitting model and thus finding a set of genes which show significant location dependent noise.

### Count data representation

A spatial transcriptomics data consists of *N* spots, with locations of the spots denoted as *X* = [*x*_1_, *x*_2_, ⋯, *x_N_*]*^T^*, where *x_i_* = [*x*_*i*_1__, *x*_*i*_2__]. Here, *x*_*i*_1__ and *x*_*i*_2__ are the horizontal and vertical coordinates of the i-th spot, respectively. We denote the relative gene expression profile (normalized and variance stabilized) for a given gene as *y* = [*y*_1_, *y*_2_, ⋯, *y_N_*]*^T^* (Figure 1(B)).

### Spatial modeling of gene expression

We assume that the expression at a particular spot depends on the location of the spot, and is also affected by the expression of nearby spots. We model this by decomposing the expression profile *y* in to two components – (1) *f*(*X*), which captures the spatial dependency of *y*, and (2) *ε*, the “noise” term that captures the part of *y* that is not explained by *f*(*X*). We use a GP prior with constant mean function *μ*(*X*) = *μ_s_* and radial basis kernel function (RBF) to model the location dependency of *y*.

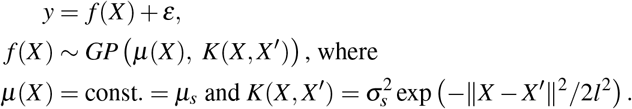

Here, *l* is called the lengthscale parameter and 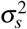 is the variance of the RBF kernel.

For the homoscedastic model, the noise term *ε* is modelled using a normal distribution with constant noise variance 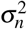,

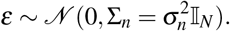

In contrast, *ε* is modelled using a normal distribution with variance at the *i*’th spot being 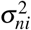 for the heteroscedasctic model,

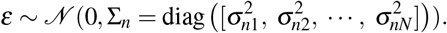

The parameters of the models, i.e., 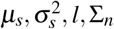, are estimated by minimizing the negative log-likelihood of *y* conditioned over the parameters. Figure 1(C) shows an example of these two models on a simulated gene expression.

### Statistical testing to detect genes with heteroscedasticity

To identify genes with significantly spatially variable noise variance, we compare the NLPD of the regular homoscedastic model NLPD_Homo_ with that of the heteroscedastic model NLPD_Hetero_. Since a lower NLPD implies a better model fitting, the genes for which the NLPD for the heteroscedastic model is significantly lower than the regular model will have location-dependent noise variance and hence denoted as *“noisy* genes” in this paper. Therefore, a statistical test on the difference between the NLPDs of homoscedastic and heteroscedastic models can be used to identify such genes. However, as the background distribution of this difference is unknown, we cannot use a z-test or t-test because they assume a normal distribution^32^. Therefore, we resort to a non-parametric test. Specifically, Wilcoxon signed-rank test^33^ was used on a set of paired NLPD values obtained by repeating the model fitting process ten times with a different random splitting of the spots into 90% training and 10% testing set (Figure 1(D)). We also used the Benjamini-Hochberg method^34^ to correct the false discovery rate (FDR).

We applied the pipeline to model gene expressions and detect *noisy* genes from two publicly available datasets — one obtained using Visium technology^5^, and the other obtained using ST technology^35^. The results, including further downstream analyses performed on the *noisy* genes to find biologically relevant signals, are discussed below.

#### NoVaTeST detects genes enriched in cancer related pathways in squamous cell carcinoma data

The first dataset contains gene expression from the skin tissue sample from a patient with squamous cell carcinoma and patient-matched normal adjacent skin sample collected using the Visium technology^36^. After QC filtering and normalization, the dataset contained normalized count data of 15725 genes expressed across 3650 spots.

Applying our model on this data, we identified 771 *noisy* genes at an FDR level of 5%. We performed enrichment analysis of the *noisy* genes to detect all statistically significant enrichment terms and hierarchically clustered them into a tree based on similarity of their gene memberships using Metascape^37^. The resulting top cluster representative enrichment terms are shown in Figures 2(A). The terms associated with cancer and immuno-response are marked bold in Figure 2(A). Interestingly, the *noisy* genes for this dataset are mostly associated with cancer-related pathways — not only cancer development^38^, tumor progression^39, 40^, angiogenesis^41^, but also cancer immuno-response^42^. This result makes sense since tumor micro-environment are highly heterogeneous due to their uncontrolled growth and thus the genes related to cancer and immuno-response are likely to have different noise variance in the tumor region compared to the tumor-adjacent healthy region.

**Figure 2.**
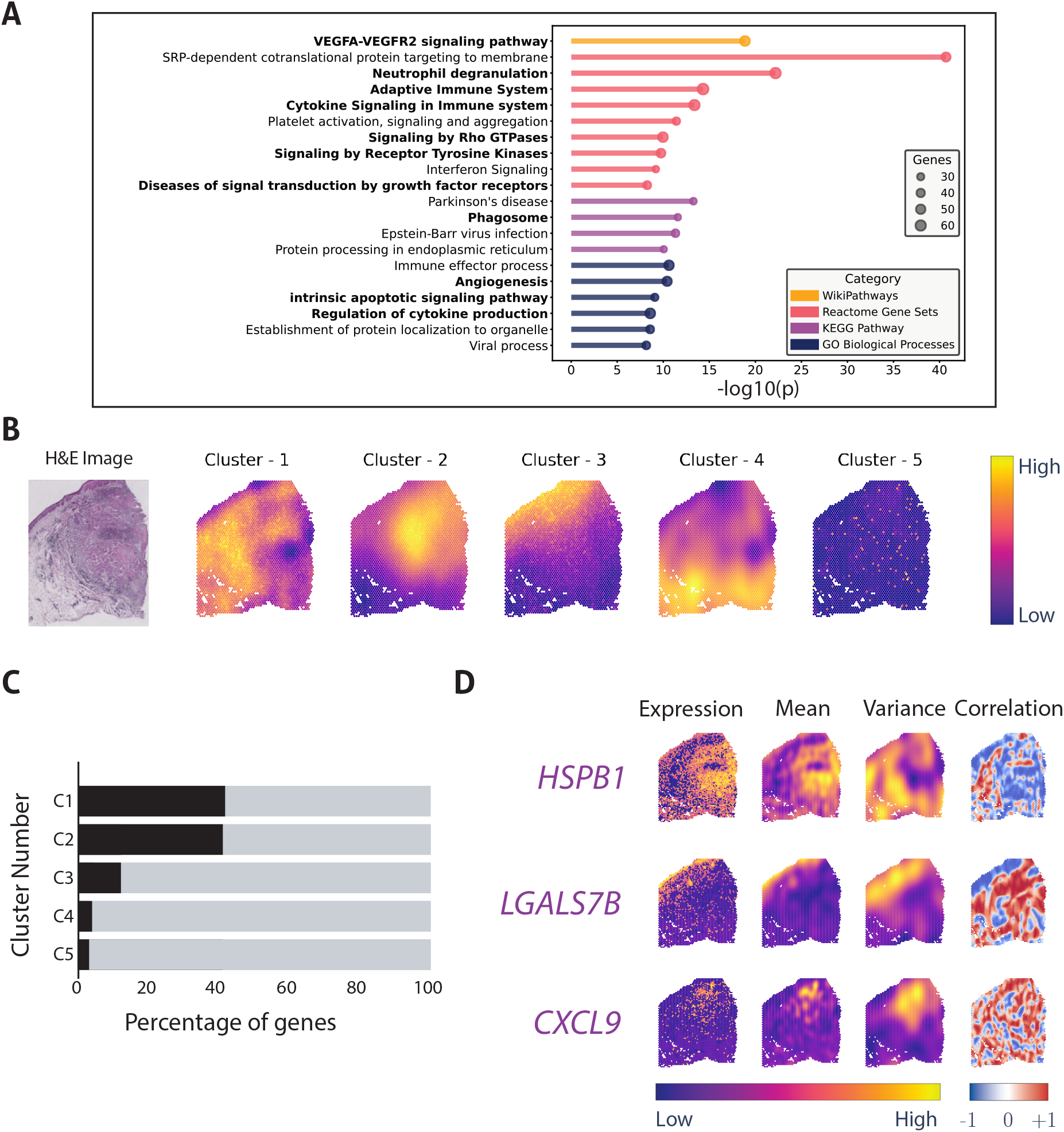
Results obtained from squamous cell carcinoma data using NoVaTeST. (**A**) Top cluster representative enriched terms for the detected *noisy* genes using Metascape. Terms associated with cancer and immuno-response are shown in bold font. (**B**) The average gene expression noise variance, averaged over the cluster members, for the five identified clusters along with the tissue H&E stained image. (**C**) Distribution of the *noisy* genes among the identified clusters. (**D**) Gene expression (log scale) and corresponding modelled mean and variance for three representative genes from the first three clusters. Also plotted is the spatial Spearman correlation between the mean and variance of the model, where the correlation for a spot is computed by considering twelve nearby spots.

The *noisy* genes were clustered based on a heuristic approach (details provided in the methods section) to find groups of genes that show visually similar noise-variance patterns. The cluster representative, which is the gene expression noise variance averaged over cluster members, for the five identified clusters in the *noisy* genes are shown in Figure 2(B). The distribution of the 771 *noisy* genes, shown in Figure 2(C), reveals that most of the genes belong to clusters 1, 2, and 3. Comparing the cluster representatives and the tissue H&E stained image, we see that for cluster 1, the variance is high mainly in the stroma and tumor-adjacent healthy region, whereas, for cluster 2, the variance is high mainly in the tumor region. For cluster 3, the variance is high along the thin line on the upper-left of the H&E image, which corresponds to the benign squamous epithelium and stroma region (see Fig. 4D of Erickson *et al*.^43^)

The log-expression and the mean and variance of the heteroscedastic model for three representative genes, *HSPB1, LGALS7B*, and *CXCL9*, selected from cluster 1, 2, and 3, respectively, are shown in Figure 2(D). *HSPB1*, which shows high mean in the tumor region and high variance primarily in the adjacent healthy and the stroma region of the H&E image, is strongly associated with tumor metastasis and metastatic colonization^44–48^. *LGALS7B* is known to play role in several types of carcinomas^49^, and the expression shows high mean in the squamous epithelium region but high variance in the stroma region. *CXCL9* from cluster 2 shows high variance in the tumor affected region and is involved in T-cell trafficking^50^. These results indicate that genes with significant spatially variable noise detected by our method indeed carry biologically significant results, and thus the importance of a generalized model.

For the selected genes mentioned above, we calculated the spatial correlation between the estimated mean and estimated variance by calculating the Spearman correlation over the twelve nearest neighbors of each spot. The results are shown in Figure 2(D). For *HSPB1*, we see that the correlation is overall low for almost all the spots. For *LGALS7B*, on the other hand, the correlation is low for the spots where the variance is high (and vice versa). These results are indication that these genes are not an artifact of the mean-variance relationship, and thus provide complimentary information to existing methods (see Discussion).

#### NoVaTeST identifies distinct noise variance patterns in cutaneous malignant melanoma data

The second dataset contains the expression of 13088 genes expressed across 279 spots from a melanoma lymph node biopsy sample collected using spatially resolved transcriptomics technology^51^. The manual annotation of the H&E image by Thrane *et. al*.^51^ shows three distinct regions — melanoma, stroma, and lymphoid (Figure 3(A)).

**Figure 3.**
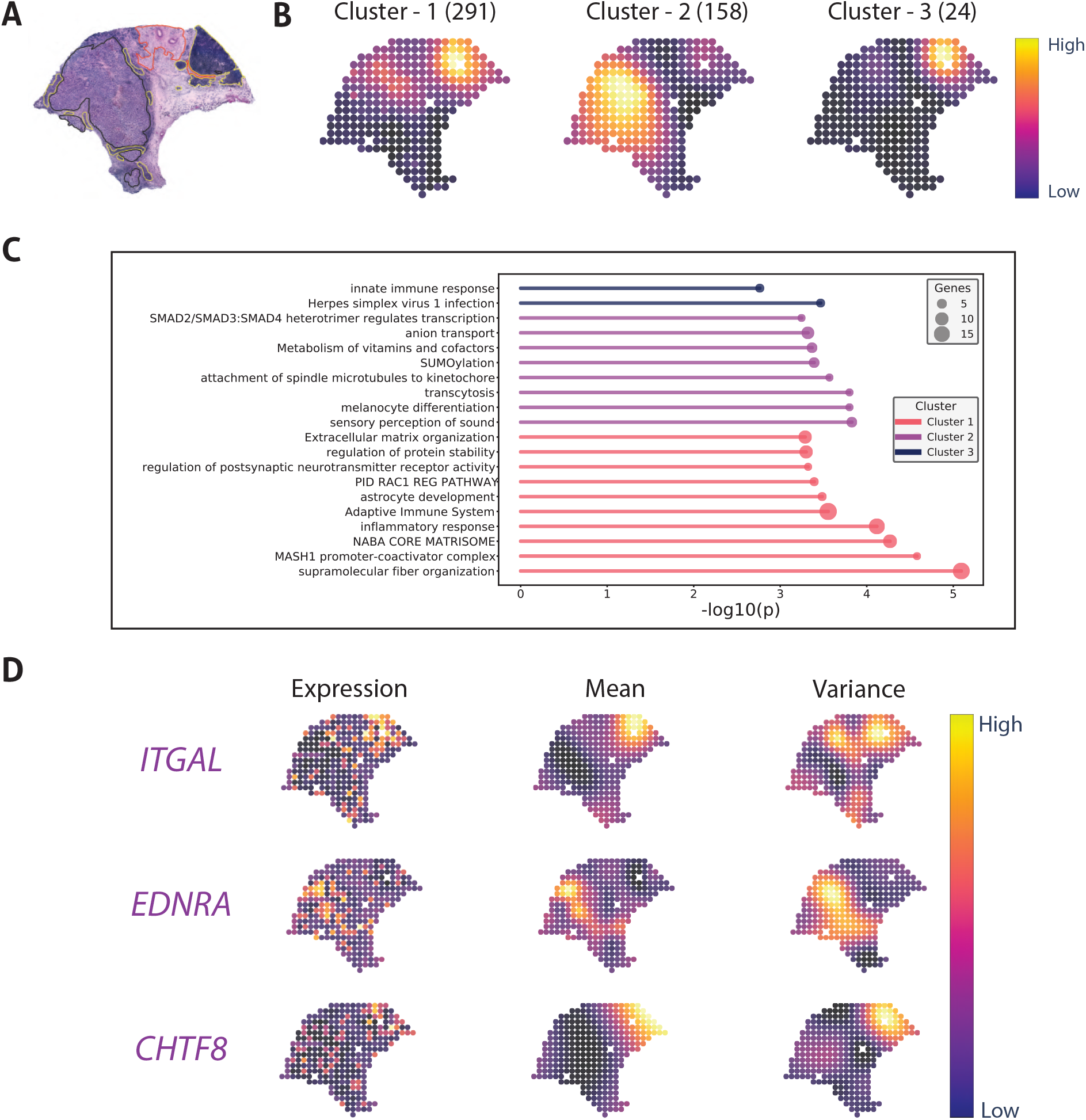
Results obtained from cutaneous malignant melanoma data using NoVaTeST. (**A**) Tissue H&E stained image of the sample with histopathological annotations - melanoma (black), lymphoid (yellow), and stroma (red), adapted from Thrane *et. al*. 2018^51^. (**B**) The average gene expression noise variance, averaged over the cluster members, for the three identified clusters. The numbers inside the parentheses denote the number of genes in each cluster. (**C**) Top cluster representative enriched terms for the detected *noisy* genes in each detected cluster using Metascape. (**D**) Gene expression (log scale) and corresponding modelled mean and variance for three representative genes from the three clusters.

Applying our model, we find 472 *noisy* genes at an FDR level of 5%. We first clustered the *noisy* genes into three clusters based on the proposed heuristic approach to elucidate the biological processes impacted by these genes. The gene expression noise variances averaged over cluster members for the three identified clusters are shown in Figure 3(B). The high variance regions of the cluster representatives of clusters 1 and 3 overlap with the manually annotated the lymphoid regions of the H&E image. On the other hand, genes in cluster 2 show high variance in melanoma region. These results point to the importance of the heteroscedastic model, as variance in phenotypically different regions show different patterns of noise variance.

Next, enrichment analysis was performed for each cluster to identify the top enriched terms. Then, the top enriched terms were hierarchically clustered based on gene memberships’ similarity. The top enriched cluster representative terms for the melanoma dataset across the three identified clusters are shown in Figure 3(C). Notably, genes in cluster 1, which show high variance in the lymphoid region, are enriched in “supramolecular fiber organization”, as well as immuno-response related terms “inflammatory response” and “adaptive immune response”. Genes in cluster 3, which also show high variance in the lymphoid region, results in only two clustered enriched terms (as there are only 24 genes), one of which is “innate immune response”. Lastly, genes in cluster 2 show high noise variance in the melanoma region, and is enriched in the GO term “melanocyte differentiation”. These results indicate that genes with similar noise-variance pattern might perform similar operations.

The log-expression and the mean and variance of the heteroscedastic model for three representative genes, *ITGAL, EDNRA*, and *CHTF8*, selected from cluster 1, 2, and 3, respectively, are shown in Figure 3(D). Abnormal expression of *ITGAL* is linked with immune regulation^52^. High expression of *ENDRA* is linked with metastasis^53^, which in this sample is also the region where the variance is high. The results indicate that the heteroscedastic model can be used to identify genes with abnormal expression in specific regions of the tissue.

## Discussion

An important first step for ST data analysis is modeling the gene expression as a function of location. There are two main types of uncertainty to consider while modeling, namely *epistemic* uncertainty and *aleatoric* uncertainty. The epistemic uncertainty is the variability of the model output due to the randomness of the model itself. In case of modeling ST data, this refers to the uncertainty (or confidence) of gene expression prediction given spatial location. On the other hand, the aleatoric uncertainty is the variability of the model output due to the randomness or noise present in the data. In this case, aleatoric variability is the part of gene expression having no spatial variation, also referred to as noise. The aleatoric uncertainty can be further classified into *homoscedastic* or input independent constant noise, and *heteroscedastic* or input dependent noise.

In this paper, we propose the NoVaTeST pipeline that uses a more generalized modeling for ST data. While existing methods use Bayesian framework to capture the epistemic uncertainty, they use a constrained assumption of homoscedastic noise to model the aleatoric uncertainty. NoVaTeST can detect the presence of heteroscedastic uncertainty, that is, genes that show significant spatial variation in noise variance.

Analysis on two different cancer ST datasets show the detected noisy genes (genes that show significant heteroscedasctic noise) in squamous cell carcinoma data are mostly associated with cancer and immuno-response related pathways. On the other hand, the noisy genes in cutaneous malignant melanoma data form three clusters in terms of noise-variance pattern, and these pattern overlap with manual annotation of different phenotypical conditions in the H&E image. These results are consistent with our initial hypothesis regarding the noisy genes.

To check whether the *noisy* genes are an artifact of mean-variance relationship, we compared the *noisy* gene list to that detected by SpatialDE^7^, a tool to detect genes with significant spatial mean expression patterns. If the detected *noisy* genes were indeed an artifact, then the *noisy* genes would also have been detected by SpatialDE. However, only about 50% of the *noisy* genes from the carcinoma data were common with the 2641 genes detected by SpatialDE (Figure S1(A)). The rest 370 *noisy* genes were not detected by SpatialDE, meaning the expression of these genes do not show any significant spatial pattern, but their noise-variance display a location-dependent pattern. For the melanoma dataset, only 32 out of the 470 *noisy* genes overlap with the 825 genes detected by SpatialDE (Figure S1(B)). These results demonstrate that the detected noisy genes are not an artifact of the mean-variance relation, rather they provide complimentary information to existing methods like SpatialDE.

Further analysis, however, reveal that the detected *noisy* genes are not completely unaffected by the mean-variance artifact, especially the *overlapping* genes detected by both NoVaTeST and SpatialDE. This can be seen from the cumulative histogram of Spearman correlation between mean and variance of the the 401 *overlapping* genes of the carcinoma dataset (Figure S1(C)). Around 50% or 200 genes show moderate correlation (between 0.5 and 0.75), meaning the mean and variance are somewhat correlated, *i.e*., spots with high expression have high variance. On top of that, around 25% or 100 genes show strong correlation between mean and variance at each spot, thus an artifact of mean-variance relation. These analyses suggest a more robust variance-stabilizing transformation should be adopted.

Gaussian processes are inherently computationally expensive. Moreover, a heteroscedastic GP, where the noise variance is modelled using another GP, does not have a closed form solution, and therefore we have to resort to iterative methods to fit a model. This further increases the runtime of the pipeline, which, again, is primarily due to computational expense of Gaussian processes. To combat this, we had to use an approximate GP where we assumed that the mean of the homoscedastic model is same as that of the heteroscedastic model.

ST data can be used to interpret cell-cell and gene-gene interactions. In the future, we want to investigate whether the noise variance pattern and the detected *noisy* genes reveal any novel information regarding cell-cell communications. Moreover, similar to recent methods like MERINGUE^54^, we want to take nonuniform cellular densities into account in our pipeline.

## Methods

### Heteroscedastic Model

A spatial transcriptomics data consists of *N* spots (one spot contains 1 to 100 cells)^6, 51^ with locations of the spots denoted as *X* = [*x*_1_, *x*_2_, ⋯, *x_N_*]*^T^*, where *x_i_* = [*x*_*i*_1__, *x*_*i*_2__] is a row matrix containing the horizontal and vertical coordinates of the *i*’th spot, respectively. We denote the relative gene expression profile for a given gene as *y* = [*y*_1_, *y*_2_, ⋯, *y_N_*]*^T^*. Due to the effect of tissue niche and inter-cellular communication, the expression at any spot is regulated by nearby spots. Therefore, the expression at a particular spot have some dependency on the location of the spot, and the covariance of nearby spots is likely to be higher. We model this by decomposing the expression profile *y* in to two components –

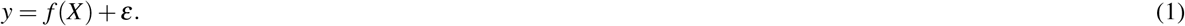

Here, *f*(*X*), which captures the spatial dependency of *y*, is a sample from a Gaussian process (GP) with mean function *μ*(*X*) = *μ_s_* and covariance kernel 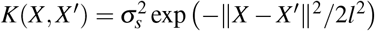, i.e., the radial basis function (RBF) kernel. The RBF kernel results in higher correlation for nearby spots as ||*X* – *X*′||^2^ is lower. The two parameters of the RBF kernel, namely the kernel variance 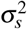 and the lengthscale parameter *l*, control the extent and smoothness of *f*(*X*), respectively. Since the number of spots is discrete, the Gaussian process boils down to a multivariate Gaussian distribution with mean vector *μ* = [*μ_s_*, *μ_s_*, ⋯, *μ_s_*]*^T^* and covariance matrix Σ*_s_*, where the (*i, j*)th entry is given by the RBF kernel, 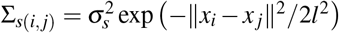. Hence, we can write

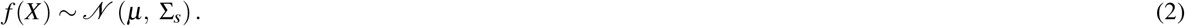

The second term, *ε*, is the “noise” term that captures the part of *y* that is not explained by the RBF kernel, which includes technical noise as well as biological noise, cell-type variation, phenotypical variation, etc. Existing tools to detect spatially variable genes assume the noise to be independent and identically distributed Gaussian. However, if the sample being analyzed is heterogeneous in terms of cell-type or phenotype (e.g., part of the sample being affected by a disease), this assumption might not hold for some genes. Therefore, we model the noise term to be a sample from a Gaussian with zero mean and covariance matrix 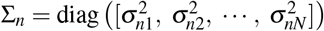, that is,

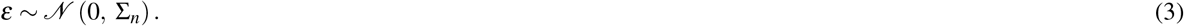

Combining equations 1, 2 and 3, we get the likelihood of y in terms of the spatial coordinates *X* and the parameters *μ_s_*, 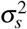, *l*, and Σ*_n_*,

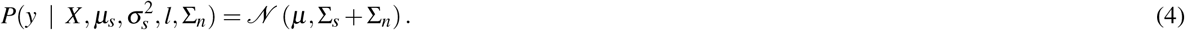

The above noise model where the noise variance is location-dependent is called a *heteroscedastic* noise, and the GP model with heteroscedastic noise is known as a heteroscedastic Gaussian Process (HGP). Usually, a second independent GP is used to model the location-dependent noise variance and thus calculate the model parameters. The posterior distribution of the HGP regression model can then be calculated by conditioning the prior HGP with the calculated parameters and training observations 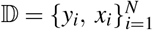. This distribution will also be a multivariate Gaussian with mean *μ_p_* and covariance Σ_*p*_ given by

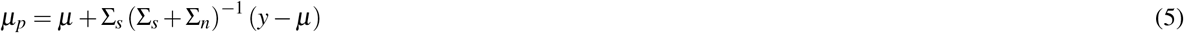

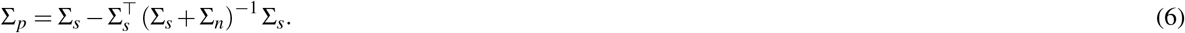

In our case, we need to model tens of thousands of genes. Hence we adopt and approximate method proposed by Urban *et al*.^55^ which provides a fast way to estimate the posterior mean and variance of the HGP model using two independent GPs. The method is briefly described below.

First, the parameters of a regular *homoscedastic* GP with 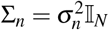 is estimated by minimizing the negative log-likelihood (from equation 4) on the given observations 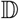,

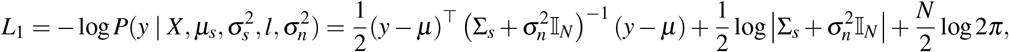

using gradient-based Broyden-Fletcher-Goldfarb-Shanno (BFGS) algorithm^56^. These parameters are then used to calculate the posterior mean *μ*_GP_ and covariance Σ_*GP*_ using equations 5 and 6. Next, a new dataset 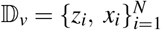 is formed, where

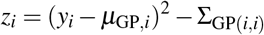

is the difference between the one-sample empirical variance at the *i* th spot and the variance for the *i* th spot. After that, a second homoscedastic GP with constant zero mean and a separate covariance matrices (∑_*sv*_ and 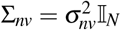) is fitted on the newly formed dataset 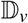 by minimizing the negative log-likelihood

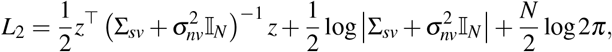

where *z* = [*z*_1_, *z*_2_, ⋯, *z_N_*]*^T^*. Finally, denoting the posterior predictive mean of the second GP regression model as *μ*_GPv_, the mean *μ*_HGP_ and variance 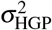 of the HGP regression model for *y* is estimated by combining the two heteroscedastic GPs according to

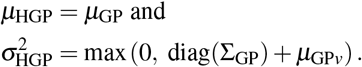

Note that for this method, both the homoscedastic GP and the (approximate) heteroscedastic GP models have same mean, and only the variance is refined for the HGP model. Finally, the negative log predictive densities on a test dataset 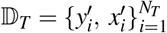 for the two models are calculated as

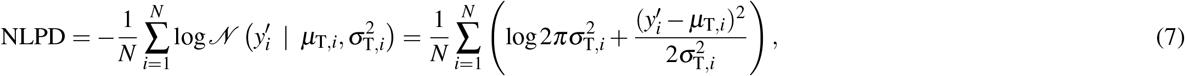

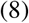

where *μ*_T,*i*_ and 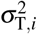 are the predicted mean and variance of the models at the *i* th spot on the test dataset 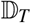.

### Data Source and QC Filtering

We have used two Spatial Transcriptomics datasets in this study. The first one is an ST data downloaded from the study of Ji *et al*.^36^ (GEO: GSE144240), which contains single cell transcriptomics data (scRNA-seq) as well as ST data from ten different patients with squamous cell carcinoma. We used the filtered ST data of patient 6 replicate 1 provided by the author (from the file `GSE144239_ST_Visium_counts.txt.gz`), which contained gene expression counts from 17736 genes expressed across 3650 spots. The second dataset was downloaded from the Spatial Research website from the study of Thrane *et al*.^51^, which contains ST data from four different patients with stage III cutaneous malignant melanoma. We used the data of patient 1 replicate 1 (from the file `ST_mel1_rep1_counts.tsv`), which contained gene expression counts from 15666 genes expressed across 279 spots. For both dataset, we filtered out the ERCC genes and mitochondrial genes, as well as practically unobservable genes that had a total count less than three. After filtering, we were left with 15733 genes across 3650 spots for the carcinoma dataset, and 13088 genes across 279 spots for the melanoma dataset.

### Clustering

The goal of the clustering algorithm is to cluster genes based on similar gene expression variance patterns. For this, we defined a heuristic distance function between the predicted variance of two genes. To calculate this distance, first the predicted noise variance for the two genes are binarized, i.e., converted to 0 or 1, by comparing the values to a threshold obtained from Otsu’s method^57^. Next, the Jaccard similarity index^58^, *J_I_*, is calculated between the binarized gene variance. Finally, the distance between the variance of the two genes is defined as *J_D_* = 1 – *J_I_*. Bottom-up (agglomerative) hierarchical clustering^59^ is performed using this *J_D_* as the distance metric. The cluster representative is calculated as the average gene expression variance (averaged over cluster members).

## Code Availability

An implementation of the NoVaTeST framework in Python is available at the GitHub (abidabrar-bracu/NoVaTeST). Notebooks used to generate the results in this paper are also available at the same repository.

