## Supplemental Figure 1 for "Identifying Genes with Location Dependent Noise Variance in Spatial Transcriptomics Data"

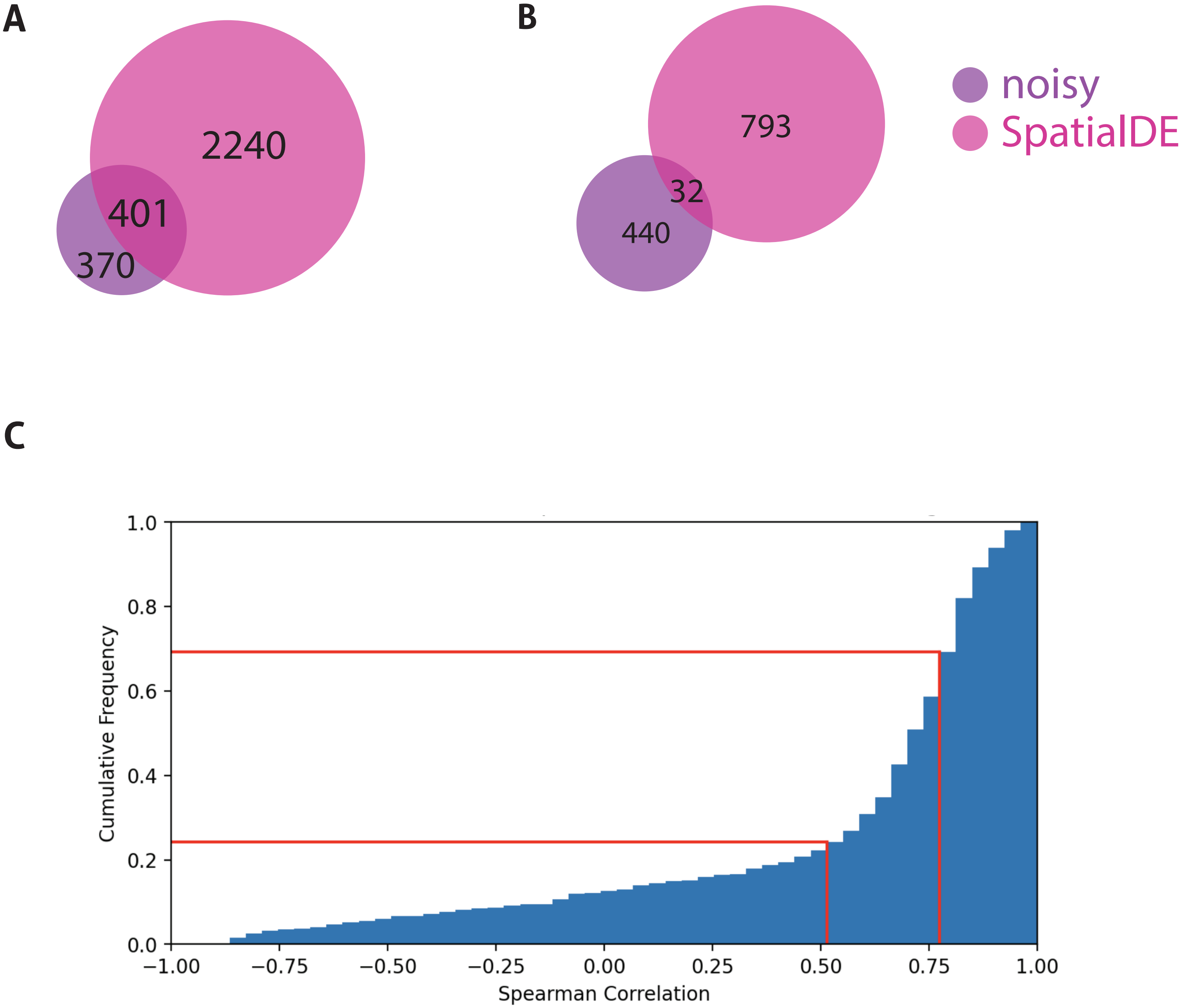


**Figure S1**. Analysis to check the existence and extent of mean-variance artifacts in the datasets. (**A**) Venn diagram showing the overlap between genes detected by SpatialDE and NoVaTeST for the carcinoma dataset. (**B**) Venn diagram showing the overlap between genes detected by SpatialDE and NoVaTeST for the melanoma dataset. (**C**) Cumulative frequency of Spearman correlation between the estimated mean and variance for the common genes (genes detected by both SpatialDE and NoVaTeST).
